# Recovery of forearm and fine digit function after chronic spinal cord injury by simultaneous blockade of inhibitory matrix CSPG production and the receptor PTPσ

**DOI:** 10.1101/2022.08.01.502398

**Authors:** Adrianna J. Milton, Daniel J. Silver, Jessica Kwok, Jacob McClellan, Philippa M. Warren, Jerry Silver

## Abstract

Spinal cord injuries, for which there are limited effective clinical treatments, result in enduring paralysis and hypoesthesia due, in part, to the inhibitory microenvironment that develops and limits regeneration/sprouting, especially during chronic stages. Recently, we discovered that targeted enzymatic modulation of the potently inhibitory chondroitin sulfate proteoglycan (CSPG) component of the extracellular and perineuronal net (PNN) matrix via Chondroitinase ABC (ChABC) can rapidly restore robust respiratory function to the previously paralyzed hemi-diaphragm after remarkably long times post-injury (up to 1.5 years) following a cervical level 2 lateral hemi-transection. Importantly, ChABC treatment at cervical level 4 in this chronic model also elicited rapid, albeit modest, improvements in upper arm function. In the present study, we sought to further optimize and elucidate the capacity for nerve sprouting and/or regeneration to restore gross as well as fine motor control of the forearm and digits at lengthy chronic stages post injury. However, instead of using ChABC, we utilized a novel and more clinically relevant systemic, non-invasive combinatorial treatment strategy designed to both reduce and overcome inhibitory CSPGs simultaneously and spatially extensively. Following a three-month upper cervical spinal hemi-lesion using adult female Sprague Dawley rats, we show that the combined treatment has a profound effect on functional recovery of the chronically paralyzed forelimb and paw, specifically during walking as well as precision movements of the digits. Our exciting pre-clinical findings will begin to enhance our understanding of the basic mechanisms underlying functionally beneficial regenerative events occurring at chronic injury stages for clinically relevant translational benefits.

**Significance statement:** Overcoming the persistent axon inhibitory environment following a functionally debilitating incomplete spinal cord lesion has long proven to be an elusive dilemma, especially months to years after the initial spinal injury. Current therapeutic and rehabilitative techniques for patients suffering from chronic cervical spinal insults minimally, if at all, address this structural hindrance and support limited return of crucial behaviors such as voluntary use of the arms and hands. Our investigation into the behavioral and anatomical consequences of systemically perturbing the high-affinity binding interaction between the receptor PTPσ and the extracellular chondroitin sulfate proteoglycans highlight an underlying barrier to the restoration of forelimb/paw walking and eating behavior 12-weeks after a cervical spinal hemi-transection.

## Introduction and Background

Each year, between 250,000 and 500,000 people worldwide suffer a debilitating spinal cord injury (SCI) with over half occurring at the cervical level, leading to millions chronically paralyzed as reported by the World Health Organization(Sekhon & Fehlings, 2001). Damage to the high cervical spinal cord can result in paralysis of much of the body, with devastating impacts on controlled, voluntary arm, hand, and digit movements leading to a severe decrease in individual autonomy and overall quality of life (Anderson, 2004; Kumar et al., 2018). The last several decades of extensive research have demonstrated the crucial role of the cell and axon growth inhibitory properties of chondroitin sulfate proteoglycans (CSPGs) during normal neural development as well as their inhibition of re-growth of cells or their processes after injury (Brittis et al., 1992; Brooks et al., 2013; Kazanis & ffrench-Constant, 2011; Lang et al., 2015; Luo et al., 2018; Snow et al., 1990; Tran, Warren, et al., 2018). In particular, we have focused on studying the role of the CSPG family of extracellular matrix molecules in the glial scar and the perineuronal net (PNN) as critical inhibitors of functional regeneration and sprouting after SCI (Alilain et al., 2011; Busch & Silver, 2007; Luo et al., 2018; Massey, 2006; Tran, Warren, et al., 2018).

Within a day after injury, CSPGs increase within and adjacent to the forming glial scar (Andrews et al., 2012; Dyck & Karimi-Abdolrezaee, 2015; Lipachev et al., 2019; D. J. Silver & Silver, 2014). This upregulation also occurs in the PNN in areas rostral and caudal to the injury site in denervated distal targets (Alilain et al., 2011; Busch & Silver, 2007; Grycz et al., 2022; Luo et al., 2018; Massey, 2006; Tran, Warren, et al., 2018). Thus, inhibitory CSPGs up-regulate expansively along the neuraxis within, proximal and distal to the lesion epicenter, leading to limited regeneration through the lesion or back into target nuclei and curtailing potential sprouting/plasticity over large distances that could occur from spared fiber systems either ipsilateral or contralateral to the lesion (Alilain et al., 2011; Andrews et al., 2012; Busch & Silver, 2007; Grimpe, 2004; Lipachev et al., 2019; Luo et al., 2018; Massey, 2006; Sakamoto et al., 2019; Tran, Warren, et al., 2018). The regulatory effects of CSPGs on both myelin and axon regeneration as well as neuronal plasticity after an acute (Dyck et al., 2015; Hartmann & Maurer, 2001; Kazanis & ffrench-Constant, 2011; Lau et al., 2013; Sherman & Back, 2008; D. J. Silver & Silver, 2014; Yamaguchi, 2000)or sub-chronic (Brittis et al., 1992; Sherman & Back, 2008; D. J. Silver & Silver, 2014; Yamaguchi, 2000)SCI, and in the context of many other conditions (Luo et al., 2018; Tran, Warren, et al., 2018) have been well documented.

The glycosaminoglycan (GAG) side chains, especially those containing CS-E, are known to bind with highest affinity to their receptor PTPσ (Miller & Hsieh-Wilson, 2015) and provide much of the inhibitory properties of CSPGs (Hussein et al., 2020; Pearson et al., 2018; H. Wang et al., 2008). Their effects can be greatly decreased by enzymatic digestion using the bacterial enzyme, Chondroitinase ABC (ChABC) (Alilain et al., 2011; Brown et al., 2012; Cafferty et al., 2008; García-Alías et al., 2009; Karimi-Abdolrezaee et al., 2012; Massey et al., 2008; Miller & Hsieh-Wilson, 2015; Ramer et al., 2014; Sakamoto et al., 2019, 2021; Sakamoto & Kadomatsu, 2017; J. Silver & Miller, 2004). ChABC is routinely used for degrading the chondroitin sulfated GAG-chains away from their resident core proteins and has been shown in various models of SCI to have pre-clinical benefits with improvements in both locomotor ability and increased axonal regeneration/sprouting toward and into previously denervated spinal sensory or motor centers (Alilain et al., 2011; Brown et al., 2012; Grycz et al., 2022; Massey, 2006; Shinozaki et al., 2016; D. Wang et al., 2011). There has only been limited success when ChABC was used alone or when coupled with other strategies, such as rehabilitative training, in sub-chronic SCI models. Somewhat improved forepaw function has been reported when ChABC was combined with pellet reaching training when the treatment was given at 4 weeks post dorsal hemisection injury (D. Wang et al., 2011). In another study, ChABC combined with lengthy treadmill training beginning 6 weeks after severe contusion cord lesion modestly improved locomotor behavior (Shinozaki et al., 2016). When ChABC was locally delivered over an extended period and combined with neural precursor cells or trophic factors, limited improvements in locomotor function have been reported in clip compression lesioned SCI animals with a 6-week delay prior to treatment (Karimi-Abdolrezaee et al., 2010). Unlike acute SCI, in the chronic state, dense glial scarring has formed around the lesion epicenter and increases in CSPGs in the PNNs has maximized within deafferented spinal levels (Andrews et al., 2012; Lang et al., 2015; Massey, 2006; Tom et al., 2004). Furthermore, the inflammatory response has largely died down and the opportunity for neuroprotection has long past (Tran, Warren, et al., 2018).

We had been focusing on the return of diaphragm function at much longer chronic stages after lateral C2 hemi-section. Recently, we documented some rather remarkable results showing complete and persistent return of diaphragm function at greatly protracted chronic stages (up to 1.5 years post lesion) upon local matrix and PNN degradation with ChABC placed directly in the vicinity of the denervated phrenic motor pool (Warren et al., 2018). The robust recovery of breathing, which occurs after a near lifetime of paralysis was surprisingly superior to anything achieved at acute or sub-chronic stages after injury using similar techniques. The ability to stimulate recovery of such strong, hemi-diaphragm function after enzyme mediated matrix degradation at C4 takes about 3 months post injury to manifest initially after hemi-lesion at C2 and this repair potential chronically, continues to grow even more over time (Warren et al., 2018). It seems likely that, in partial injury models, improvements via enzyme administration in past studies may have been limited, at least in part, because not enough time had been allowed to pass prior to treatment to enable the slow process of potentially meaningful spontaneous plasticity to reach sufficient levels. While able to breathe with the intact hemi-diaphragm, ambulate, feed, and drink, high cervical hemisected animals also exhibit long-lasting deficits in forelimb behavior, including reduced gross and fine movements of the ipsi-lesional forelimb and the paw. Our previous ChABC enzyme applications at C4 to the very long chronically C2 injured animal, in order to restore diaphragm function, were not optimized for effects upon the forelimb, which is innervated from spinal levels around C5-C8 (Warren et al., 2018; Warren & Alilain, 2019). However, the strategy, none-the-less, elicited rapid albeit relatively small positive changes in upper arm function, likely mediated via enzyme diffusion caudally towards the cervical enlargement (Warren et al., 2018).

The use of ChABC has not been applied clinically due to a variety of potential complications such as heat lability of a bacterial enzyme which, in addition, must be administered intraparenchymally into the injured spinal cord, which can further traumatize an already damaged CNS structure. Further, the injection field is limited to a discrete area, so positive behavioral changes are dependent upon the precision of locally restricted alterations in the matrix. Several labs have engineered stable, longer lived ChABC or more widespread viral vector release strategies although the direct delivery issue remains with these approaches (Bartus et al., 2014; Burnside et al., 2018; Hettiaratchi et al., 2019, 2020; Lee et al., 2010). In order to target CSPG mediated inhibition more globally, we have focused our attention on the possibility of using systemic agents that could potentially diminish the abundance of CSPGs or disrupt the association with their major receptor without directly touching the spinal cord in the presence of any evolving lesion. To accomplish this, we utilized a high dose of a subcutaneously delivered protein tyrosine phosphatase receptor (PTPσ) inhibitor, intracellular sigma peptide (ISP). This TAT-wedge domain peptide has been shown to work acutely in vitro and in vivo to prevent the conversion of growth cones into a dystrophic state through the excessively tight substrate adhesion mediated via the PTPσ receptor and the CSPGs (Fisher et al., 2011; Ham et al., 2019; Sakamoto et al., 2019; Shen et al., 2009). In addition, we investigated the therapeutic effectiveness of a clinically approved, orally administered small molecule perineuronal net inhibitor, 4-methylumbelliferone (we call it PNNi for use in the CNS), which serves to limit a major transmembrane scaffold for PNN assembly resulting in the PNN unable to be stably formed or maintained (K.-A. Irvine et al., 2014). Thus, in addition to the use of the receptor disrupting wedge peptide (ISP), we employed a novel strategy in combination which can simultaneously and expansively reduce the matrix PNN CSPGs (K.-A. Irvine et al., 2014).

Our chronic LC2H SCI model allows us to test the role of PNN associated CSPGs in curtailing potential functional sprouting from the intact side of the cord at the level of the cervical enlargement or likely elsewhere within the CNS related to forearm and paw function. By strongly interrupting the PTPσ-CSPG interaction we were able to significantly restore functional movements of the impaired forelimb during over ground locomotor behavior as well as cereal manipulations by the fingers during eating. The synergistic effect of ISP and PNNi further supports the crucial role of CSPG mediated inhibition via the PTPσ receptor in curtailing functional synaptic plasticity at lengthy chronic time-points following SCI.

## Methods

### Ethical declaration and animal husbandry

All experiments were approved by the Institutional Care and Use Committee at Case Western Reserve University (CWRU), Cleveland. Animals were housed in groups of two or three, exposed to a normal dark–light cycle with free access to food, water, and environmental enrichment ad libitum. The health and welfare of the animals was monitored daily by the study investigators and veterinary staff at Case Western Reserve University.

### Cervical level two hemitransection (LC2H) injury and systemic treatment(s)

Spinal cord injury surgeries were performed as previously described (Warren et al., 2018). Adult female Sprague Dawley rats (280 ± 20 g; Harlan Laboratories Inc., Indianapolis, IN, USA) were anaesthetized with an intraperitoneal injection of ketamine+xylazine cocktail (70 mg kg^−1^/7 mg kg^−1^). The dorsal neck-shoulder area was shaved, cleaned, and sanitized using Betadine and 70% ethyl alcohol, and analgesics were administered through subcutaneous injection of meloxicam (1 mg kg^−1^). Body temperature was maintained and monitored throughout the surgery at 37 ± 1 °C. A dorsal midline incision ∼3 cm in length was made over the cervical region. After the skin and paravertebral muscles were retracted, a laminectomy was performed over C2 and C3 and the rostral spinal cord was exposed. A 21G syringe needle was positioned at the midline of the spinal cord, and a left lateral durotomy and hemisection were performed caudal to the C2 dorsal roots making certain that the needle tip extended to and was dragged along the ventral boney lamina surface. This process was repeated five times and ranged from the midline to the most lateral extent of the spinal cord. The muscle layers were sutured together with 3-0 Vicryl and the skin closed using wound clips. The animals were given meloxicam (1 mg kg^−1^) and sterile saline subcutaneously for up to 5 days post-surgery along with nutritional support should the animals’ weight have dropped more than 5% of that pre-injury. The anatomical completeness of the injury was confirmed through microscopy and behavioral assessment. While animals were able to survive immediately post-injury and exhibited no signs of malnutrition or infection, animals that exhibited persistent autophagy as a result of the injury before or during the systemic treatment phase were excluded from the study. Prior to the initiation of the systemic treatment(s), all animals were confirmed to achieve a performance score no better than 4.5 ± 1 out of 17 at three days post-injury as described in the locomotor assessment, and 8 ± 1 twelve weeks post-injury (Fig. 1; see *Methods-Forelimb Locomotor Scale* (Singh et al., 2014)). For cereal eating assessments, all animals were confirmed to receive a performance score no better than 0 ± 1 out of 9 at three days post-injury, and 2 ± 1 twelve weeks post-injury (Fig. 3; see *Methods-Irvine, Beattie, and Bresnahan Forelimb Recovery Scale* Irvine et al., 2014).

**Figure 1.**
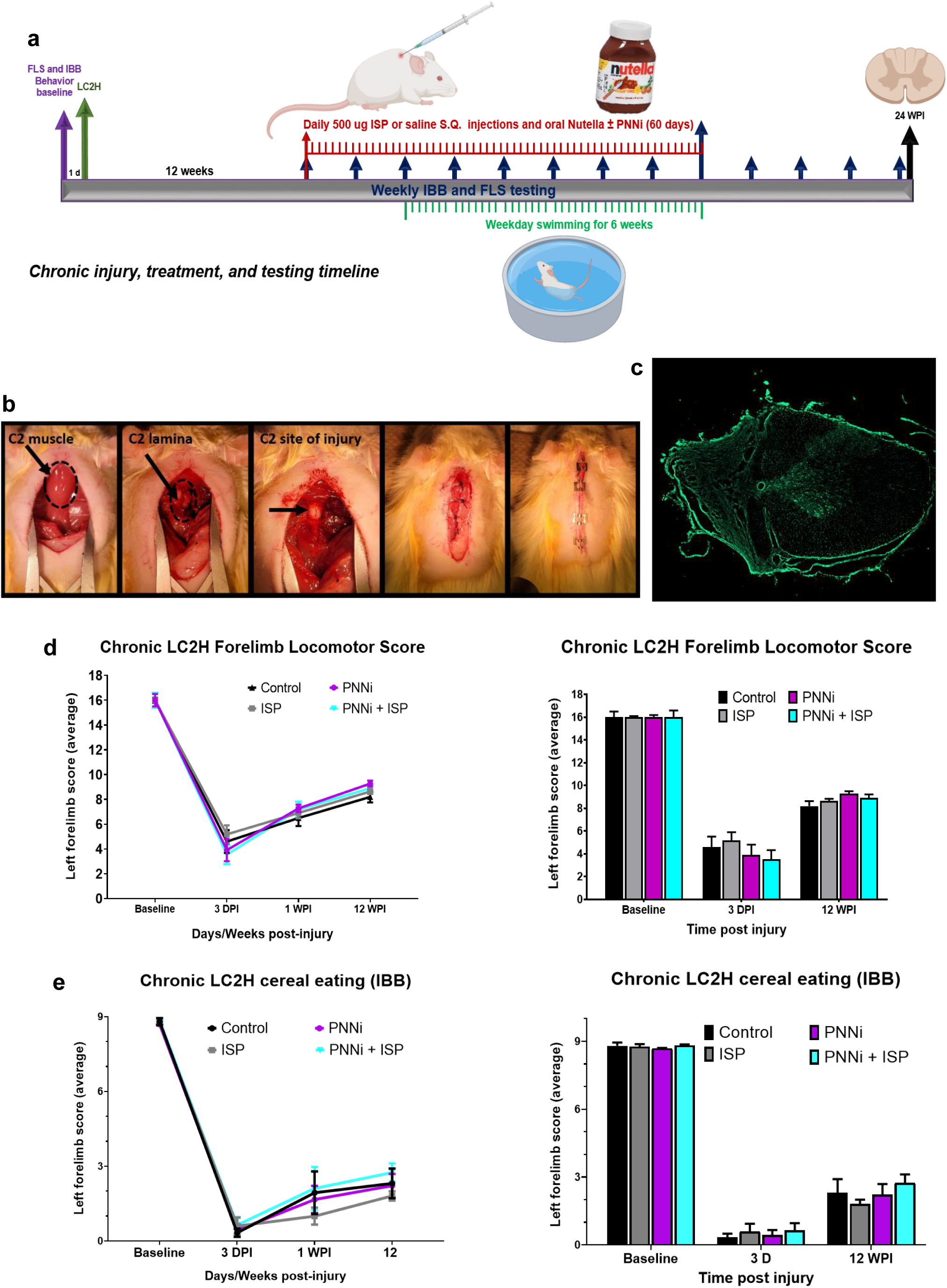
A-D Incomplete cervical SCI severely impairs forelimb function during walking and eating behavior at chronic stages. (A) Timeline of the experimental protocol. (B) Representative image depicting surgical SCI procedure at C2. All animals received a left hemi-lesion at C2 removing ipsilateral descending motor inputs. (C) Representative image of injured spinal cord at C2 (scale bar = 500 um). (D and E) LC2H severely impairs forelimb walking ability and cereal eating behavior at 3 days, 1 WPI, and 12 WPI. There were no significant differences across the experimental treatment groups compared to control.

Three months after LC2H, the rats began a systemic treatment protocol that included a daily 0.5 mL subcutaneous injection of 500 ug of intracellular sigma peptide (ISP), combined with oral feeding of Nutella mixed with the small molecule perineuronal net CSPG inhibitor, PNNi. Control animals were injected with saline and fed Nutella not containing PNNi. Two separate cohorts received ISP or PNNi alone. These systemic treatments were administered with the goal to simultaneously disrupt the interaction between would-be sprouting axons and the PTPσ receptor as well as to reduce the density of inhibitory extracellular perineuronal net components. Each cohort (n = 8 – 10) received the treatments daily for 60 days and was tested blindly and weekly for arm and hand function for the duration of treatment and continuing four weeks following the termination of treatments. Our assessments (see below for details) of forelimb/paw function were conducted using the forelimb locomotor scale (FLS) (Singh et al., 2014) and the IBB forelimb recovery scale (K.-A. Irvine et al., 2014). Both are rating scales similar to the well-known BBB scoring method for overground walking (Basso et al., 1995) but are focused on the forelimb and digits.

### Behavioral training and assessments

Animals were acclimated to the laboratory room testing environment and cereals in the home cage for five consecutive days. The animals were also handled by the researchers twice a day to minimize any stress/anxiety prior to exposure to the testing apparatuses measured by the animal’s reliability to perform the assessment. The following week, the animals were acclimated to the various testing apparatuses which included a ten-minute session each day in the glass cylinder used in the cereal eating assessment. The animals were trained to eat at least 3 of each cereal type (sphere-shaped i.e. Cocoa Puffs, or donut-shaped i.e. Fruit Loops) within the ten-minute session. For acclimation to the FLS testing platform, rats were placed on top of an open field platform approximately 4’ in diameter and allowed to explore for 5 minutes. FLS and IBB pre-training were conducted for five consecutive days. Baseline behavior performance was acquired following training and before LC2H (Fig. 1A).

### Forelimb function assessments

Forelimb function was determined through assessments which involved monitoring behavior that the animals performed naturally. This included the forelimb locomotor assessment scale (FLS) (Singh et al., 2014) and the Irvine, Beattie and Breshnahan (IBB) forelimb recovery scale (K.-A. Irvine et al., 2014). Baseline values were taken prior to injury and at 3 months after injury just prior to daily subcutaneous injection of high dose ISP or saline alone or combined with Nutella ± PNNi administration (for 60 days) and then weekly following treatment application (Fig. 1A). Statistical comparisons were made between treatment groups and within behavioral measurements using two-way, repeated-measures ANOVA with post-hoc Bonferroni Correction (Graphpad Prism).

Forelimb Locomotor Scale (FLS): These testing methods were established previously (Singh et al., 2014). Briefly, rats are encouraged to continuously walk on top of an elevated circular platform approximately 4’ in diameter. Rats that remained stationary for longer than 10-15 sec were enticed to move by having them follow a pencil or a piece of paper, or by lightly tapping or scratching on the side of the open field. If the animal failed to respond to these stimuli, it was picked up by the forequarters and placed in the center of the open field or opposite its previous position, which usually caused it to move. Rats were recorded during locomotion using a high-speed (60 FPS) camera for offline scoring to assess forelimb walking functionality. Videos were scored in a blinded fashion in slow-motion (at least 50%) playback using the Lightworks video editing software. Recorded videos containing 3-4 minutes of the animal freely walking on the elevated platform were measured using an 18-point scale (0-17). Animal locomotor behaviors were assessed for impairments and assigned a score reflecting the ability of the animal to perform steps that were consistently plantar, parallel, and weight-bearing (Singh et al., 2014).

The Irvine, Beattie, and Bresnahan (IBB) Forelimb Recovery Scale: These testing methods were established previously (K.-A. Irvine et al., 2014). Briefly, animals were placed in the 10” H x 8” W clear glass cylinder that was used for previous acclimation and training. Each rat was given one sphere-shaped (Cocoa Puff) and one donut-shaped (Fruit Loop) cereal to eat. Their manipulations of the cereals were recorded using a high-speed (60 FPS) camera for offline scoring to measure forelimb and digit abilities during eating. Videos were scored blinded in slow-motion (at least 50%) playback using the Lightworks video editing software. Recorded videos containing the animal eating one sphere-shaped cereal (i.e. Cocoa Puff) and one donut-shaped cereal (Fruit Loop) were measured using a 10-point scale (0-9) (K.-A. Irvine et al., 2014). The animal’s ability to dexterously manipulate the cereals was assessed for impairments and assigned a score reflecting the ability of the animal to grasp the cereal with subtle adjustments made using the second, third and fourth digits and wrist movements (K.-A. Irvine et al., 2014).

### Stimulation of forelimb utilization

All animals were placed in a pool (4 feet diameter) filled to a depth of ∼18 inches at a temperature of 35-38 degrees Celsius to match the rat’s natural internal temperature. Animals were placed in the pool for 1 minute. 2-4 rats were placed in the pool at one time. After swimming, rats were partially dried with a cotton towel and then placed in empty cages lined with paper towels and allowed to further dry themselves and groom uninterrupted for 1 hour at room temperature. The rats were then replaced back into their home cages. Swimming took place 5 days a week beginning two weeks after treatment application began, excluding the day prior to and the day of behavioral acquisition using the FLS and IBB assessments described above (Fig. 1A).

### Histology for serotonergic fibers

At 20 WPI, animals were anesthetized with a cocktail of ketamine and xylazine prior to cardiac perfusion with 4% paraformaldehyde dissolved in PBS. Spinal tissue containing regions of interest were dissected and post-fixed in 4% paraformaldehyde overnight. Prior to sectioning, the tissue was cryoprotected with sequential treatments first in a solution of 30% sucrose in PBS and then in a 1:1 mixture of 30% sucrose in PBS + OCT. After embedded in OCT, the tissue was sectioned to 30 μm coronal samples and free-floating sections were prepared for subsequent immunofluorescence analysis using standard protocols. Primary antibodies targeting SERT (1/1000; EnCor Biotechnology; Cat No. RPCA-SERT) and WFA-Lectin (1/1000; Vector Laboratories; Cat No. B-1355) were coupled to Alexa Fluor 488 and Alexa Flour 555 conjugated secondary antibodies (1/500), respectively. Nuclei were visualized using Hoechst 33342 (1/3000; Life Technologies; Cat No. C10338). All stained sections were mounted onto slides, coverslipped using an antifadent mounting medium, and examined using an inverted Leica SP8 confocal microscope.

### Statistical assessment

#### Histology for serotonergic fibers, perineuronal net, and incomplete cervical injury

At 20 WPI, animals were anesthetized with a cocktail of ketamine and xylazine prior to cardiac perfusion with 4% paraformaldehyde dissolved in PBS. Spinal tissue containing regions of interest were dissected and post-fixed in 4% paraformaldehyde overnight. Prior to sectioning, the tissue was cryoprotected with sequential treatments first in a solution of 30% sucrose in PBS and then in a 1:1 mixture of 30% sucrose in PBS + OCT. After embedded in OCT, the tissue was sectioned coronally at 30 μm and free-floating sections were prepared for subsequent immunofluorescence analysis using standard protocols. Primary antibodies targeting SERT (1/1000; EnCor Biotechnology; Cat No. RPCA-SERT) and WFA-Lectin (1/1000; Vector Laboratories; Cat No. B-1355) were coupled to Alexa Fluor 488 and Alexa Flour 555 conjugated secondary antibodies (1/500), respectively. Nuclei were visualized using Hoechst 33342 (1/3000; Life Technologies; Cat No. C10338). All stained sections were mounted onto slides, coverslipped using an antifadent mounting medium, and examined using an inverted Leica SP8 confocal microscope or Zeiss Axio Imager microscope.

For lesion area and volume, sixteen spinal cords were selected for analysis from the total available using a random number generator. Sections were obtained using a cryostat at 30 μm/section spanning a ∼15 mm long cervical segment containing the LC2H. Spinal cord lesion area and volume were analyzed through fluorescent Nissl staining (ThermoFischer) performed on cross-sectioned tissue of each spinal cord at 500-μm intervals. The coverage of the lesion as well as the location was determined using a Zeiss Axio Imager microscope and quantified blindly using NIH ImageJ software analysis of 3-4 sections per animal by using monochromatic image and background readings subtracted at the defined area of interest. In all subjects, the left half of the cervical spinal cord was completely severed from the midline to the lateral most extent of the tissue and there were no deviations (Fig. 1c).

## Results

### All treatments improve gross function following chronic cervical hemi-SCI

All rats displayed a baseline pre-injury FLS performance level at a score of 17, or consistent plantar stepping of the forepaw, predominant parallel placement and continuous toe clearing (Singh et al., 2014) (Fig. 1D and Fig. 2A). At 3 days and at 1 week post-injury (WPI), all rats exhibited an FLS score early on of ∼4 and a bit later on improving slightly to ∼7, respectively (Fig. 1D and Fig. 1A {3 days: control = 4.6 ± 0.9; ISP = 5.1 ± 0.7; PNNi = 3.9 ± 0.9; ISP+PNNi = 3.5 ± 0.7} {1 WPI: control = 6.5 ± 0.6; ISP = 6.9 ± 0.4; PNNi = 7.2 ± 0.3; ISP+PNNi = 7.1 ± 0.6}). These findings demonstrate the consistency of the targeted removal of descending supraspinal motor inputs upon inducing the lateral transection across all of the treatment groups immediately following SCI. Unsurprisingly, by 12 WPI, all rats had reached a pre-treatment baseline FLS score of ∼8 indicating that the animals displayed some spontaneous recovery of function that was consistent prior to treatment and not statistically different between groups (Fig. 1D and Fig. 2 {12 WPI: control = 8.2 ± 0.4; ISP = 8.6 ± 0.2; PNNi = 9.2 ± 0.2; ISP+PNNi = 8.9 ± 0.3}). An FLS score of 8 denotes the animals’ ability to perform partial weight-supported steps but only on the dorsal surface of the impaired limb/paw (Fig. 1D and Fig. 2) (Singh et al., 2014).

**Figure 2.**
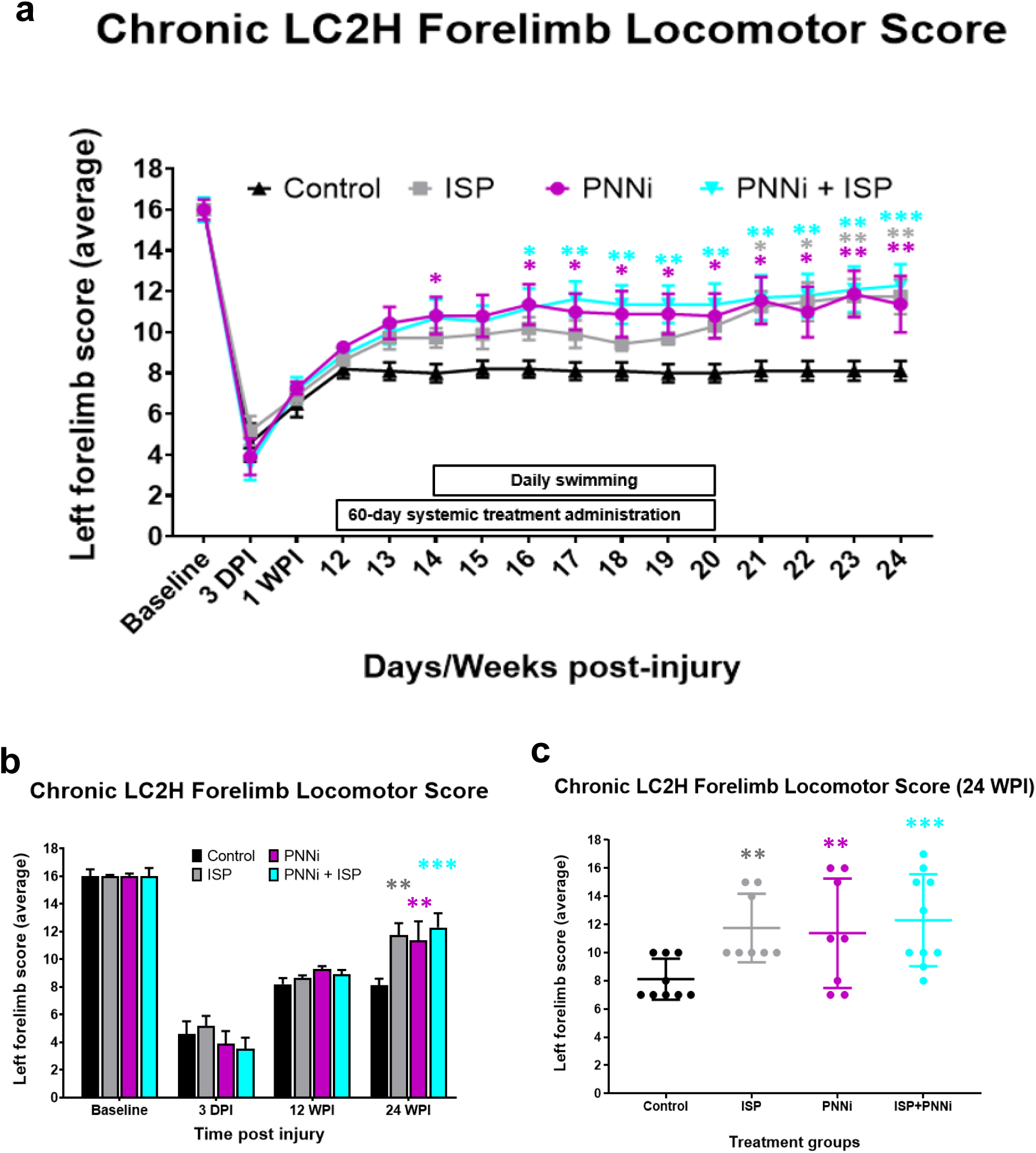
A-C The Forelimb Locomotor Score is significantly improved following systemic treatment to remove CSPGs in the PNN and/or modulate receptor PTPσ. (A) Chronic SCI rats were measured for forelimb locomotor function before and after LC2H and systemic ISP ± PNNi. Both ISP ± PNNi significantly improved weight-bearing stepping after chronic PTPσ-PNN binding perturbation. (B – C) At 24 WPI, animals receiving either systemic drug showed an average improvement in walking behavior, with combinatory recovery showing the greatest significance and rate of recovery.

Interestingly, the control group never gained further FLS assayed function beyond that demonstrated in the pre-treatment assessment, and therefore, any recovery observed is likely due to the administered treatment (Fig. 2B-C and Fig. 3 {12 WPI: 8.2 ± 0.4 v 24 WPI: 8.1 ± 0.4}). The various treatments caused a fairly rapid increase in performance as early as the first week after their onset (Fig. 2 {13 WPI: control = 8.1 ± 0.43; ISP = 9.7 ± 0.55; PNNi = 10.4 ± 0.77; ISP+PNNi = 10 ± 0.5} {ANOVA, ISP p = 0.678, F = 1.627; PNNi p = 0.132, F = 2.355; ISP+PNNi p = 0.4712, F = 1.9}). There was a significant difference between the FLS score at 14-WPI in the PNNi alone treated animals (Fig. 2A {ANOVA, p = 0.037, F = 2.818}), while the combination treatment group revealed a significance in recovery beginning at 16 WPI which then improved slowly during the drug administration phase (Fig. 2A {ANOVA, p = 0.0279, F = 2.982}). At 24 WPI, the control rats maintained an average FLS score of ∼8, while rats that received ISP alone, PNNi alone, or in combination displayed significant improvements in forelimb recovery during walking compared to the control group (Fig. 2 and Fig 3 {ANOVA, ISP p = 0.0089, F = 3.639; PNNi p = 0.0261, F = 3.264; ISP+PNNi p = 0.0007, F = 4.189). There was evidence of synergy between the 2 treatments. Thus, behavioral recovery occurred especially in rats that received the combined treatment of ISP and PNNi. At 24 WPI, ISP alone and PNNi alone treated animals reached an average FLS score of ∼11, which functionally translates to continuous plantar stepping but with poor wrist control and no toe clearance (ISP = ± 0.86; PNNi = 11.3 ± 1.37). The combination treated rats received an average final score of ∼12, indicating continuous weight bearing and plantar stepping that is predominantly parallel but without toe clearance. However, the best responding animals in the group receiving both PNNi and ISP reached strikingly high performance levels (Fig 1D and Fig. 3C {highest 24 WPI FLS: ISP+PNNi = 17}).

**Figure 3.**
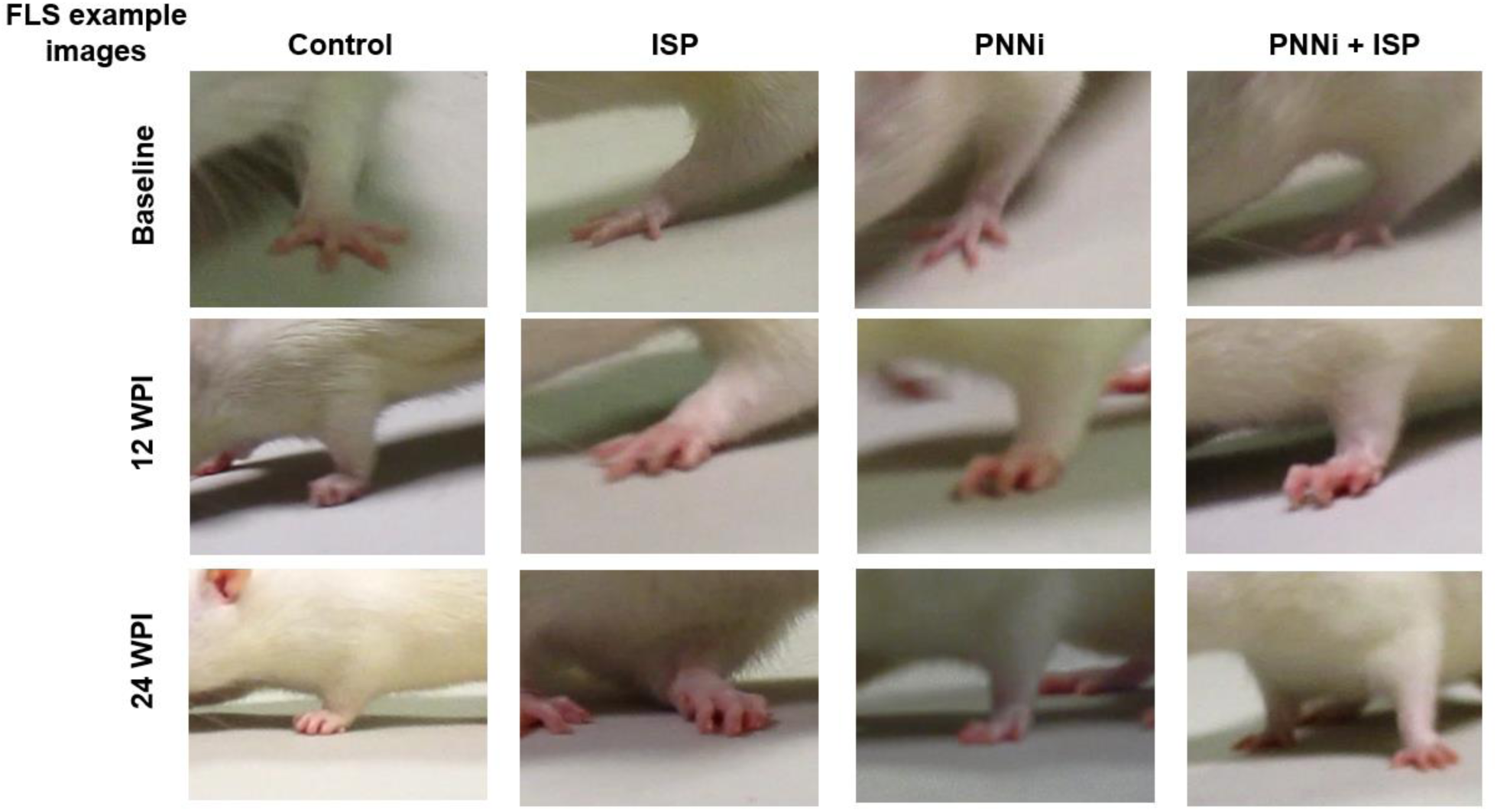
Forelimb locomotor scale walking assessment. Example still-frame images from actively walking rats at baseline, 12WPI, and 24 WPI (four weeks after systemic treatment ended) during FLS analysis (\**P* < 0.05, *\*\*P <* 0.005, *\*\*\*P* < 0.0005) *refer to* Figure 2.

### The combination treatment best improves skilled forelimb and digit function following a partial chronic cervical SCI

When assessing for cereal eating ability, all rats displayed a baseline pre-injury performance level at a score of 9 when both the soon to be injured (left) and uninjured (right) were measured. Normal performance is described as almost always maintaining properly shaped paws (conforming to that of the cereal) during grasping of the cereal with fully flexed elbow, extensive distal limb movements, subtle cereal adjustments and volar supported manipulations (K.-A. Irvine et al., 2014). At all weekly behavioral testing time-points before and during systemic treatment administration, the uninjured forelimb displayed consistent normal function during cereal eating. At 3 days (DPI) and at 1 week post-injury (WPI), all rats exhibited an IBB score of ∼0.5 and ∼2, respectively (Fig. 1E and 4A-B {average IBB score 3 DPI: control = 0.3 ± 0.1; ISP = 0.6 ± 0.3; PNNi = 0.4 ± 0.2; ISP+PNNi = 0.6 ± 0.3} {average IBB score 1 WPI: control = 1.9 ± 0.8; ISP = 1.0 ± 0.3; PNNi = 1.6 ± 0.5; ISP+PNNi = 2.1 ± 0.8}). By 12 WPI, all rats reached a pre-treatment baseline IBB score of ∼2, which translates to cereal eating behavior that uses the non-volar surface of the injured forelimb, with a predominant fixed forepaw position that is clubbed into a fist-like flexed state, while some animals display an extended forepaw phenotype (K.-A. Irvine et al., 2014) (Fig. 1E, Fig. 4A-B and Fig. 5). Unlike the FLS data detailed above, the IBB assessment did not reveal a significant effect of treatment until week four of treatment administration. Furthermore, the significant behavioral recovery observed was only demonstrated in the animals that received both PNNi and ISP in combination (Fig. 4A, Fig. 5; average IBB score 17 WPI: control = 2.5 ± 0.6; ISP+PNNi = 4.8 ± 0.5, {ANOVA, *p* = 0.014, *F* = 2.30}).

**Figure 4.**
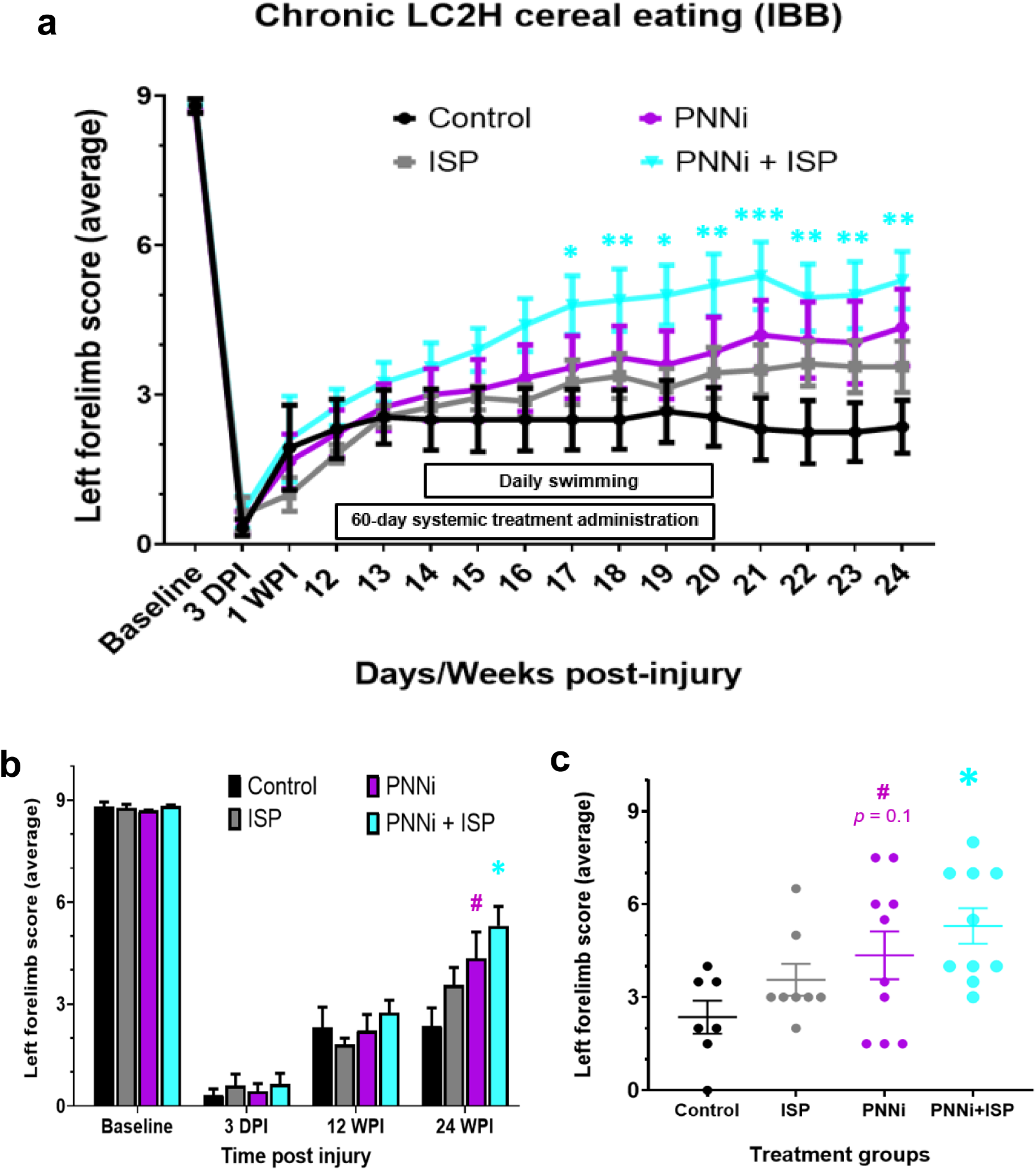
A-C Cereal eating ability assessed using the IBB-rating scale is significantly improved following combined systemic treatment to both remove CSPGs in the PNN and modulate receptor PTPσ. (A) Chronic SCI rats were measured for forepaw cereal manipulability before and after LC2H and systemic ISP± PNNi. Only animals receiving combinatory therapy showed significant improvement as early as 17 WPI. (B – C) Only chronic injured animals receiving ISP + PNNi improved significantly in cereal eating behavior by 24 WPI. Rats that received PNNi alone demonstrated a trend towards significance 4 weeks after the termination of oral administration.

**Figure 5.**
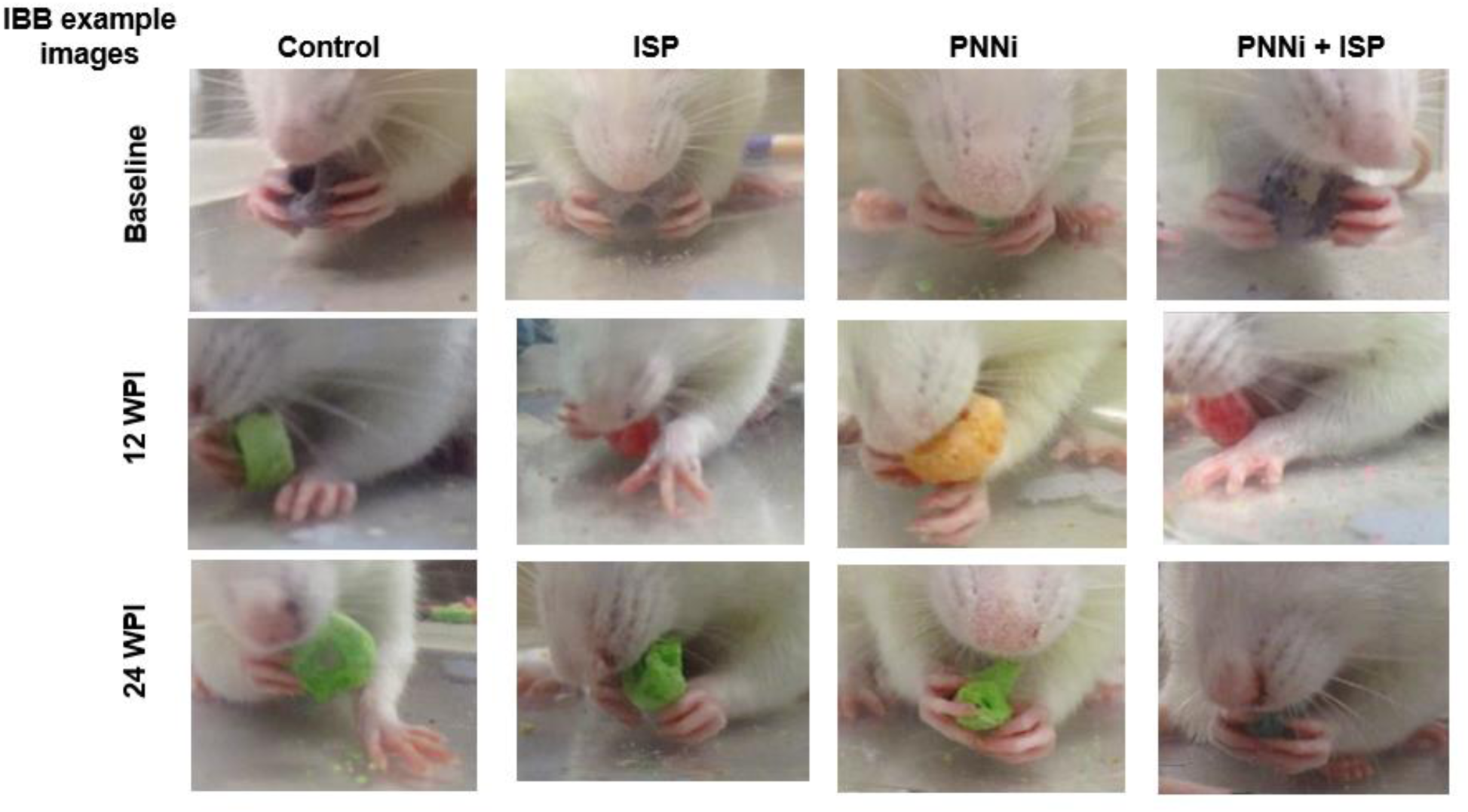
IBB cereal eating assessment of fine digit and wrist functional recovery. Example still-frame images of animals eating cereal at baseline, 12WPI, and 24 WPI (four weeks after systemic treatment ended) during **IBB** analysis (\**P* < 0.05, *\*\*P <* 0.005, *\*\*\*P* < 0.0005) *refer to* Figure 4.

Our behavioral results using the IBB assessment to measure forelimb and paw cereal manipulations further demonstrated that the combination treatment group eventually developed the ability to use their once paralyzed paw/fingers remarkably better than controls. The control animals recovered paw/finger function only minimally over time (compared to normal animals) to an IBB score between 2 and 3, where the affected paw may remain partially clubbed and mostly rests on the ground and where, at best, some crude, exaggerated cereal adjustments are made (K.-A. Irvine et al., 2014) (Fig. 4C and Fig. 5, average IBB score 24 WPI: control = 2.3 ± 0.5). The paw does not exhibit adaptability and fails to conform to the shape of the piece of cereal. While animals receiving either ISP or PNNi alone did not show a similar effect of meaningful forelimb functional recovery as we showed in the FLS assessment, these animals did show a strong trend towards significance by the end of the study compared to the control group (Fig. 4 and Fig. 5; {average IBB score 24 WPI: ISP = 3.5 ± 0.5; PNNi = 4.3 ± 0.7} {ANOVA, ISP *p* = 0.5, *F* = 1.20; PNNi *p* = 0.07, *F* = 1.99}). On the contrary, the combination of ISP and PNNi led to clear improvements in paw/finger function during cereal eating (K.-A. Irvine et al., 2014). On average, at 24 weeks post injury and with the combinatorial treatment, the once paralyzed animals recovered to an IBB score of between 5 and 6 with extensive contact manipulatory movements (K.-A. Irvine et al., 2014) (Fig. 4 and Fig. 5; average IBB score 24 WPI: ISP+PNNi = 5.3 ± 0.6; {ANOVA, *p* = 0.02, *F* = 2.94}). Our most encouraging finding was that the best responding animals, especially when eating a donut-shaped cereal such as a Fruit Loop, could achieve a remarkable score of 7 (9 is the highest possible score) (K.-A. Irvine et al., 2014) where sometimes normal, shape adapting grasping can occur. In summary, these behavioral data confirm the hypothesis that modulating CSPGs in the scar and especially in the PNN as well as their receptor simultaneously is central to the recovery of precision digit function long after SCI.

### Histological analyses reveal dense sprouting of 5-HT axons in the spinal cord of combination treated animals

In our previous studies showing the therapeutic efficacy of ISP **acutely** after severe contusive spinal cord injury (Lang et al., 2015), we demonstrated extensive sprouting of serotonergic axons at spinal levels below the lesion. Upon examination of 5-HT axons just caudal to the chronic injury site in our combination treated animals, we have again observed this 5-HT axonal sprouting phenomenon (Fig. 6). At the end of our behavioral study (24 WPI), we visualized CSPG immunoreactivity using Wisteria floribunda agglutinin (WFA) staining, focusing on the ventral horn grey matter at spinal levels caudal to the LC2H near the forelimb muscle innervating motor pools for all experimental groups. In control rats that were chronically injured at C2 and received no systemic treatment, there was an upregulation in the accumulation of CSPGs in the extracellular matrix (ECM) compartment and in PNNs surrounding cells in the cervical enlargement compared to animals that received ISP with or without oral PNNi (Fig. 6). This increased CSPG signal was observed throughout the entire cervical region (C5-C8) and was seen on both the injured and uninjured sides suggesting, as have others (refs), that much of the spinal column distal to the lesion responds to a very localized (C2) SCI by upregulating the production of CSPGs over a wide distance.

**Figure 6.**
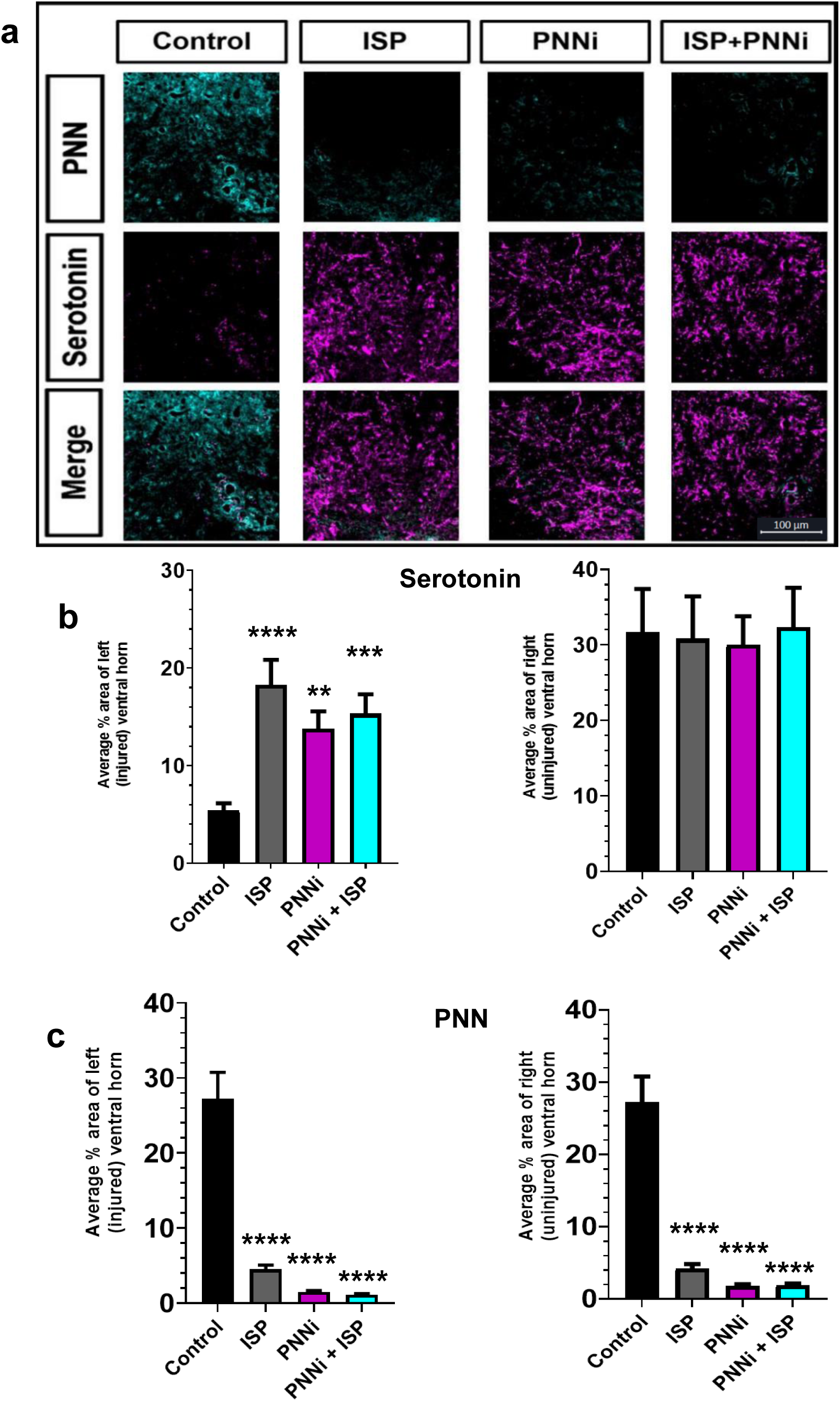
A-C PNNs are decreased and serotonergic fibers are significantly increased in the cervical enlargement grey matter. (A) Example immunofluorescent image of densely sprouted ventral horn serotonergic fibers (magenta) immediately caudal to the LC2H in an animal treated with combined ISP and PNNi. (B-C) Quanitifications of analyzed images shown in (A). Serotonergic fibers are significantly increased, while PNNs demonstrate a drastic inversely correlated decrease within the extracellular grey matter of the cervical enlargement. These results are not observed when chronic cervical SCI rats receive saline + Nutella as control.

Additionally, the increased CSPG intensity persists for months (24 WPI) after the initial spinal insult has occurred (Fig. 6). Therefore, in the chronic stage of SCI, CSPGs within the ECM and PNNs are ideal targets for restoration of forelimb movements. Along with the increase in the CSPG matrix there was an obvious decrease in the presence of 5-HT+ fibers within the cervical ventral horn grey matter (Fig. 6). Together, these data suggest that the observed behavioral deficits following chronic LC2H are preceded by increased PNN-CSPGs and loss of descending motor input, including serotonergic axon innervation onto downstream targets. We then compared the CSPG and 5-HT+ fiber densities in animals that received ISP ± PNNi to the control animals described above. In animals receiving a daily systemic treatment to either inhibit the PTPσ receptor alone (ISP) or block the production and secretion of hyaluronan (PNNi) alone, a significant decrease of CSPGs in the ECM and PNNs on the injured (left) and uninjured side of the spinal cord was observed (Fig. 6; ANOVA, ISP ipsilateral p < 0.0001, contralateral p < 0.0001; PNNi ipsilateral p < 0.0001, contralateral p < 0.0001). A similar (but not beyond that which occurred in the single treatment animals) reduction in extracellular CSPGs happened again when both systemic treatments were administered together during the chronic SCI phase (Fig. 6; ANOVA, ISP+PNNi ipsilateral p < 0.0001, contralateral p < 0.0001).

In addition, we measured the serotonergic axon innervation density in the same region. We found that animals that received a systemic treatment daily, either alone or in combination, exhibited a significant increase in 5HT+ fibers within the cervical enlargement compared to animals with unperturbed PNNs (Fig. 6; ANOVA, ISP ipsilateral p < 0.0001; PNNi ipsilateral p < 0.0033; ISP+PNNi ipsilateral p < 0.0004). Not surprisingly, we did not observe a difference in the serotonergic innervation on the contralateral, uninjured side (Fig. 6; ANOVA, ISP contralateral p = 0.991; PNNi contralateral p = 0.9898; ISP+PNNi contralateral p = 0.9993). These findings are consistent with previous studies that have successfully stimulated serotonin axon plasticity following SCI (Li et al., 2015; Rink et al., 2018; Tran, Sundar, et al., 2018; Warren et al., 2018). Descending serotonergic fibers are known to influence, indirectly or directly, the excitability of spinal cord motoneurons and it is well established that serotonin (5-HT) plays an important role in modulating locomotion (Jacobs & Fornal, 1999; Sakai et al., 2000; Wei et al., 2014). Many studies have shown that functional recovery following a SCI is potently influenced by increased numbers of serotonin axons at spinal levels caudal to the initial injury1,4– 6(Lang et al., 2015; Tran, Sundar, et al., 2018; Warren et al., 2018). Taken together, these findings support our hypothesis that interrupting the PTPσ-CSPG binding interaction promotes axon sprouting at spinal levels caudal to the partial cervical SCI and may help underly our observed restoration of forelimb walking. Whether serotonin plays a role in restoring fine digit control is not well undertood, but there are several studies that have shown a role for serotonin in the modulation of certain aspects of flexor (Seo et al., 2011; Vitrac & Benoit-Marand, 2017).

## Discussion

We assessed functional recovery of forelimb behavior in a hemisection model of chronic cervical SCI and have revealed, for the first time, that treatments with PNNi and ISP alone but especially in combination can stimulate a good measure of recovery. Furthermore, these functional improvements in forelimb movement persisted for at least 4 weeks after the daily treatment(s) had ended. When given for two months following a chronic LC2H, ISP or PNNi administered alone were able to improve proximal and distal forelimb behavior during more crude locomotion. Moreover, we assessed fine motor behaviors during cereal eating and showed rather remarkably improved digit function when we administered our systemic treatments together.

It is quite interesting that recovery of forearm function begins to occur relatively rapidly following systemic treatment in our chronic injury model. Surprisingly, behavioral improvements occur much more quickly than they do after acute injury^14^. This timeframe mimics that which we have shown previously in our ChABC treated chronic LC2H SCI respiratory model suggesting, again, that sprouting likely had already occurred but is somehow being masked by the CSPG component of the PNN (Warren et al., 2018). Thus, in our 12-week cervical hemi-lesion model, certain sub-populations of the contralateral descending supraspinal tracts with small numbers of axons that re-decussate within the spinal cord caudal to the level of injury (Bareyre et al., 2004), or intervening interneurons with similar properties of plasticity (Fenrich & Rose, 2009), may be very slowly sprouting new processes that re-cross the midline to the denervated hemisphere ipsilateral to the lesion even in the presence of the PNN (Alilain & Goshgarian, 2008; Friedli et al., 2015; Hawthorne et al., 2011; Lane et al., 2008; Porter, 1895). Various motor systems underlying forelimb and paw movement, including the serotonergic, propriospinal, and cortico-spinal neurons have an innate propensity to sprout on their own, although at a very diminished rate (Asboth et al., 2018; Bareyre et al., 2004; Cafferty et al., 2008; Fenrich & Rose, 2009; García-Alías et al., 2015; Hawthorne et al., 2011; Ishida et al., 2019). These new collaterals could serve as latent, or inactive synapses which over time could be beneficial, harmless or possibly even disadvantageous to recovery (Ueno et al., 2016; Warren et al., 2018). Our results demonstrate that the potential activity of such sprouting can be supportive to functional recovery but is largely kept dormant by CSPGs within the PNN in the vicinity of the relevant motor pools.

What is the mechanism by which the PNN may curtail synaptic activity and perhaps help to create silent or latent synapses? Increases in PNN density and changes in the chondroitin sulfation pattern to a more growth inhibitory state occurs during normal aging (Foscarin et al., 2017) and after spinal cord (Carulli & Verhaagen, 2021; Lipachev et al., 2019). It is well established that the PNN as well as the LAR family of CSPG receptors play important roles not only in regulating various morphological aspects of axonal sprouting during the development of critical periods (Dyck & Karimi-Abdolrezaee, 2015; Lipachev et al., 2019; Snow et al., 1990) as well as after injury in the adult (Andrews et al., 2012; Brown et al., 2012; Goussev et al., 2003; Hussein et al., 2020; Lang et al., 2015; Massey et al., 2008; D. J. Silver & Silver, 2014; Tom et al., 2004) but also in the physiological control of synaptic activity (Carceller et al., 2022; Carulli & Verhaagen, 2021; Gottschling et al., 2019; John et al., 2022; Sclip & Südhof, 2020). Interestingly, in a model of Alzheimer’s disease, the PNN in the cortex encroaches into synaptic clefts which shrinks the perimeter of ECM barriers around the PNN holes, thereby markedly reducing the diameter of the portals for communication between pre and post-synaptic elements (Stoyanov et al., 2021). However, the invasion of the net in terms of its potent function smothering effect on the synapse has not been elucidated (Lewis & Brookhart, 1951). Latent synapses are known to occur throughout the CNS (Atwood, 1999). One of the most interesting and well-known latent descending motor projections is the crossed phrenic pathway (Alilain & Goshgarian, 2008; Goshgarian, 2003, 2009; Lewis & Brookhart, 1951). It is already present but can be re-awakened in the adult in seconds after an LC2H injury by creating anoxic stress via lesion of the contralateral phrenic nerve (Alilain & Goshgarian, 2008; Warren et al., 2018). Perhaps, in addition to other proposed mechanisms that can lead to synaptic latency, synapses of sprouted axons in the cervical enlargement could be made inactive via this same type of matrix reorganizing phenomenon (Buttry & Goshgarian, 2014; Chen et al., 2018; Goshgarian, 2003, 2009; Lewis & Brookhart, 1951; Weng et al., 2017).

Recently, an independent study confirmed the functional benefits of subcutaneous peptide treatment showing that 500 ug/day of ISP given acutely after severe contusive injury greatly improved hindlimb locomotor, sensory and bladder recovery (Rink et al., 2018). Indeed, with the use of the well characterized BBB locomotor rating scale, their contusion-injured animals (that stabilized at a control baseline score of 6) on average recovered by a remarkable 9 full points to a level of 15 (Rink et al., 2018). Importantly, while bladder (but not walking) function showed a dose response when given ISP at a maximum of 44 ug/day (Lang et al., 2015), the much higher dose of 500 ug/day provides the first evidence for a dose response by ISP in promoting enhancements in locomotor behavior, at least after acute SCI. Similarly, ISP has some therapeutic effects after acute T8 cord hemisection in the adult rat when assessed by an automated gait analyses system (Ham et al., 2019, 2020). ISP has also now been shown to be effective in promoting regeneration and functional recovery after SCI when delivered via a plasma exosome based biological scaffold (Ran et al., 2022) as well as a self-assembling peptide hydrogel (Sun et al., 2023) The regenerative potential of blocking the PTPs receptor by ISP has, in addition, been demonstrated in models of MS (Luo et al., 2018), ischemic and hemorrhagic stroke (Luo et al., 2022; Yao et al., 2022), ischemic heart attack (Gardner et al., 2015; Sepe et al., 2022), and peripheral nerve injury (Li et al., 2015; Lv & Wu, 2021). The high dose of 500 ug/day of ISP is that which we chose to use and was effective in our chronic study. However, the most optimal dosing regimen, the best route of delivery as well as the optimal amount of peptide that are most functionally beneficial have not yet been established.

Hyaluronan (HA) is the major scaffold for PNN assembly and if HA synthesis is blocked then the PNN is not able to structure itself properly (Duncan et al., 2019; Nagy et al., 2015; Rink et al., 2018). In addition to the use of the wedge peptide ISP, we employed a novel strategy in combination which can substantially reduce the PNN CSPGs within the CNS via systemic delivery of an already clinically approved small molecule proteoglycan synthesis inhibitor called 4-Methylumbelliferone (4MU) or hymecromone (Caon et al., 2020; Duncan et al., 2019; Nagy et al., 2015). Referred to as PNNi in our study, we repurposed this small molecule in our model of chronic SCI. One of the major challenges for translation of SCI regenerative strategies from rodent to human is the enormous relative increase in size of the human which necessitates careful considerations of concentrations, toxicity and routes of delivery to optimize the amount and time of residence of the therapeutic drug within the CNS. Oral delivery of a small molecule inhibitor that is already clinically approved is a preferred strategy for drug delivery. A novel therapeutic approach to overcome the inhibitory effects of the PNN involves curtailing the production of HA. PNNi specifically inhibits HA synthesis by acting as a competitive substrate for hyaluronan synthase, an enzyme critical for HA production (Nagy et al., 2015) Preliminary findings have shown that inhibiting HA synthesis with PNNi resulted in significant decreases in the hyaluronan link protein as well as the amount of CSPGs in an in vitro model as well as in the brain and spinal cord. In vivo, the reduction of the CSPG component of the PNN was especially prominent in the ventral horn around motor neurons (S. F. Irvine et al., 2023; Štepánková et al., 2023). We therefore hypothesized that PNNi, and especially when administered together with ISP, could be an obvious dual therapy since the combined strategy would decrease the ligand as well as block the receptor globally and could be an effective, clinically viable treatment method following chronic partial cervical lesions. Indeed, we confirmed that chronic disruption of proper PNN assembly in the 2 cohorts of animals that received 4-MU (+/-ISP) significantly reduced the presence of WFA+ CSPGs within the cervical enlargement. Importantly, the reduction in CSPGs was inversely correlated with the increased density of serotonergic axons observed within the ventral grey matter of the same region. PNNi as well as ISP are non-invasive treatments with low toxicity that can be stopped at any time to reverse the receptor blockade and HA-synthesis inhibiting effects of the combination and allow for the PNN to be re-established. This can help avoid potential long term adverse side effects and help stabilize newly formed synapses.

While reduction of CSPGs in the 4-MU treated animals was expected, why was there also a reduction of CSPG matrix, again where 5-HT axon densities had increased, in the ISP only treated animals? A variety of motile cell types, including neurons, use tightly regulated release of proteases to remodel the surrounding extracellular matrix along their potential routes of migration (Even-Ram & Yamada, 2005; Krystosek & Seeds, 1984; Tran, Sundar, et al., 2018; Zuo et al., 1998). We have described, in vitro and in vivo, an interesting downstream event in both DRGs and serotonergic neurons that occurs as a consequence of RPTP σ modulation by ISP resulting in the locally enhanced release of the matrix degrading enzyme, cathepsin B. Cathepsin B is a proteolytic enzyme that is normally produced, albeit in limited amounts, by growing neurons during crucial periods of extracellular matrix remodeling and is also augmented in neurons within the spinal cord grey matter after injury (Ellis et al., 2005; Turk et al., 2012). The exaggerated secretion of Cathepsin B, likely at the leading edge of ISP-treated axon growth cones, helps them to navigate past or within an inhibitory CSPG barrier (Ellis et al., 2005; Tran, Sundar, et al., 2018; Turk et al., 2012). We have shown that this phenomenon of enzyme hyper-secretion is protease specific and occurs not only in neurons but also in multiple migratory cell types that bear the PTP receptor as they encounter CSPGs. Thus, in oligodendrocyte progenitor cells in models of MS as well as SVZ derived neural progenitor cells in an ischemic model of malignant stroke, ISP specifically induced MMP2. The protease encouraged digestion of glial scar associated CSPGs which, in turn, restored stem cell migrations and their differentiation resulting in functional improvements (Luo et al., 2018, 2022; Tran, Sundar, et al., 2018).

Our behavioral findings demonstrate a robust return of forelimb function months after a partial spinal transection during gross and fine voluntary motor behaviors that are innate and ecologically relevant to our model organism. The ability to recover these behaviors is more than likely owed to sprouting of their underlying neural pathways, particularly serotonergic, corticospinal or propriospinal tracts descending contralateral to the SCI and forming latent, re-decussated terminals within the spinal cord cervical enlargement containing forelimb muscle motor neurons. Thus, we were able to induce plasticity of the spinal tissue by manipulating the presence and effect of the PNN on spared, sprouting descending axons using treatments that either inhibited the PNN-binding neuronal receptor RPTPσ (via ISP), or impaired the PNN to be synthesized and maintained. These results suggest that the systemic targeting of the PNN and PTPσ receptor at chronic SCI periods is a viable therapeutic target for treatment that can be extended or ended at any time or combined with other beneficial interventions such as rehabilitative training or neuromodulation, with limited adverse side-effects.

## Acknowledgments

We thank Dr. S. Fischer and Dr. K Brock for their dedicated veterinary assistance. We further thank Dr. W. Alilain, and also Dr. P. Popovich, Dr. D. McTigue, and Dr. M. Basso (the OSU Spinal Cord Injury Course) for surgical and behavioral technical training. Financial support was provided by Wings for Life, the Ohio Department of Higher Education Third Frontier Program, The Brumagin-Nelson Fund, The Griffin Family, The Timothy Brodigan Trust, The Kaneko Family Fund and NIH NINDS grants (1R01NS101105 and 1R011NS113831). We acknowledge the support of the Government of Canada’s New Frontiers in Research Fund (Mend the Gap), NFRFT-2020-00238.

